# The human GABA_A_ receptor α4 subunit variant Ile114Asn linked to epilepsy impairs membrane expression with β3 and δ subunits

**DOI:** 10.64898/2026.04.22.720185

**Authors:** Draegan Watson, Cecilia M. Borghese, Kristina Jülich, Audrey C. Brumback, Marcel P. Goldschen-Ohm

## Abstract

GABA_A_ receptors mediate fast inhibitory neurotransmission in the brain, and variants in their subunits are linked to neurological disorders like epilepsy. For many identified variants, it remains to be determined whether the variant is pathogenic. Here, we functionally characterized the α4 subunit variant p.I114N (p.I79N in the mature protein), found in an individual with epilepsy, using patch-clamp electrophysiology in combination with β3 and δ subunits in HEK293T cells. Whereas the variant has little to no impact on basal activity or GABA-sensitivity, lower maximal GABA-elicited currents as compared to wild-type channels suggest that it impairs assembly or trafficking to reduce expression of channels in the plasma membrane.

## Description

Epilepsy is a complex neurological disorder that often arises from genetic variants affecting key components of neural signaling, such as GABA_A_ receptors. These receptors are critical for mediating fast inhibitory neurotransmission in the brain, and disruptions in their function can lead to abnormal neuronal activity and seizures (Absalom et al., 2024; Hernandez & Macdonald, 2019). Despite advances in available treatments, many individuals with epilepsy due to suspected genetic variants experience limited therapeutic success (Orsini et al., 2018), as current interventions may not address the underlying molecular mechanisms. Complicating this challenge is the uncertainty surrounding the pathogenicity of numerous identified GABA_A_ receptor variants, which hinders accurate diagnosis of the underlying cause of disease and the development of personalized treatment strategies. Therefore, there is an urgent need to systematically determine the functional impact of these potential pathogenic variants to improve outcomes for affected individuals.

The GABA_A_ receptor α4 subunit variant I114N (I79N using the mature protein numeration, see Reagents), was identified in a six-year-old boy with a developmental and epileptic encephalopathy characterized by medically refractory seizures with global developmental delay and developmental regression. Whole exome sequencing revealed a heterozygous missense variant of uncertain significance (VUS) in *GABRA4* c.341T>A p.I114N. This variant is located at the β3/α4 inter-subunit interface in the extracellular domain above the GABA binding site (Fig. 1A), where it could potentially disrupt either subunit association or GABA binding. Given that other variants in *GABRA4* have been associated with neurological phenotypes such as developmental delay, behavioral abnormalities and epilepsy (Ahring et al., 2022; Sajan et al., 2024; Vogel et al., 2022), we determined the impact of the I114N variant on basic channel function to assess its potential pathogenicity.

**Figure 1.**
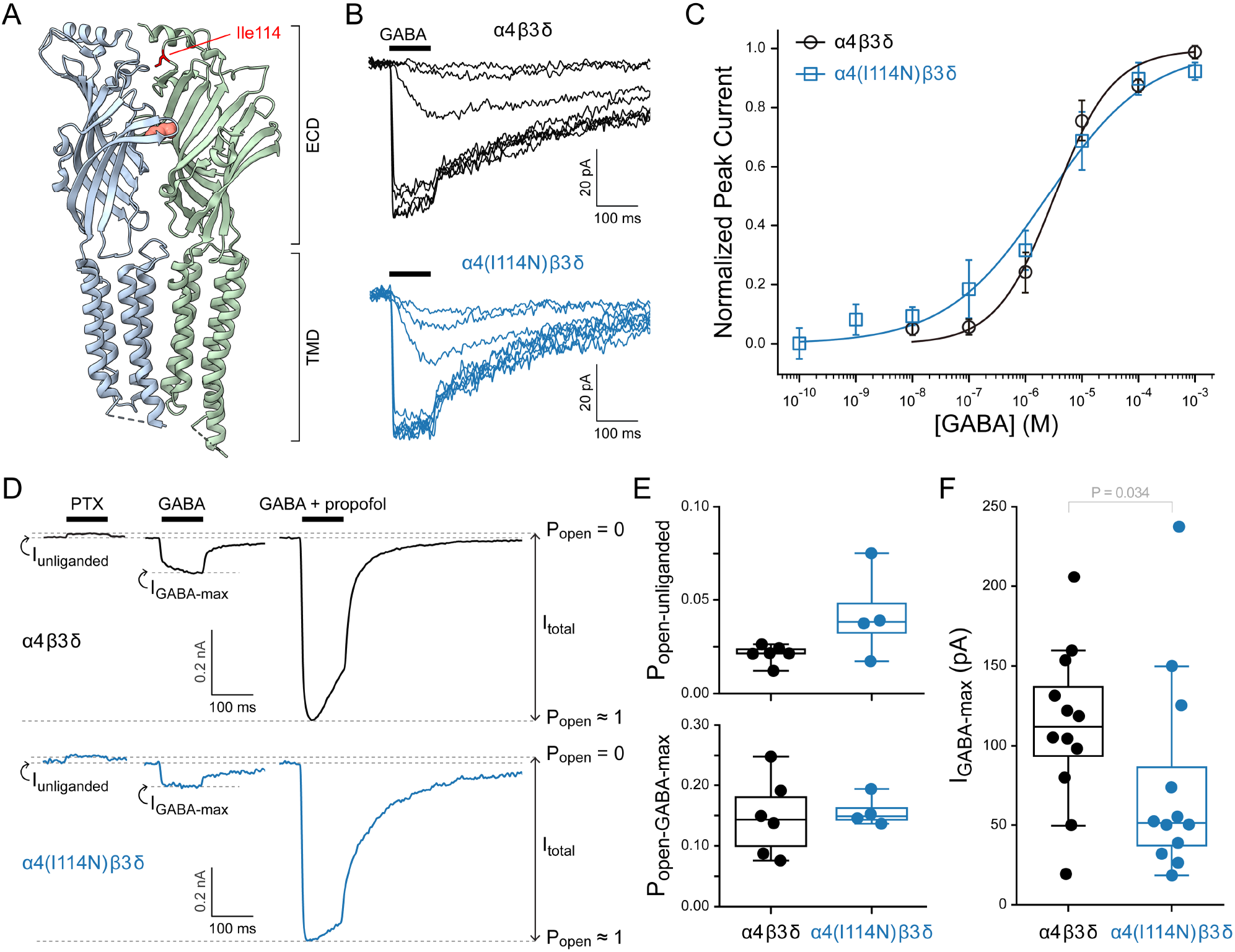
The GABA_A_ receptor variant α4(I114N) has normal basal and GABA-elicited function when expressed in HEK-293T cells with β3 and δ subunits. (**A**) CryoEM map (PDB 7QN7) for a α4β3δ GABA_A_ receptor. Only two of the five subunits are shown with extracellular (ECD) and transmembrane (TMD) domains indicated. Residue Ile114 (red sticks) is located above the GABA (salmon spheres) binding site in the ECD. Visualization with ChimeraX (Meng et al., 2023). (**B**) Family of current responses to a series of 100 ms pulses (black bar) of GABA from low nanomolar to 1 mM concentration. (**C**) Summary of normalized GABA concentration-response curves across cells as illustrated in panel A. Data points are mean ± sem, and solid lines are fits to the combined data across cells with the Hill equation (Eq. 1). Fit parameters (EC_50_, *h*) and number of cells (n) are α4β3δ: EC_50_ = 4.3 μM, *h* = 0.9, n = 5; α4(I114N)β3δ: EC_50_ = 2.4 μM, *h* = 0.5, n = 4. (**D**) Current responses in the same cell to either 1 mM PTX, 1 mM GABA, or the combination of 1 mM GABA and 30 μM propofol. Open probability (P_open_) was estimated by normalizing from the zero-current baseline in the presence of pore blocker PTX to the current amplitude elicited with the combination of GABA and propofol. This illustrates that GABA alone is relatively inefficient at opening the channel, whereas addition of propofol enables more robust activation. (**E**) Summary of basal unliganded and GABA-elicited open probabilities as illustrated in panel D. Data points are individual cells. Box plots indicate median and quartiles. Number of cells (*n*) are α4β3δ: n = 6; α4(I114N)β3δ: n = 4. No differences based on t-Test (P_open-unliganded_: p = 0.18; P_open-GABA-max_: p = 0.34). (**F**) Summary of peak GABA-elicited currents from all experiments as illustrated in panels B and D. Data points are individual cells. Box plots indicate median and quartiles. Number of cells (*n*) are α4β3δ: n = 12; α4(I114N)β3δ: n = 12. Statistical significance was assessed with a Kolmogorov-Smirnov t-Test (p = 0.033) due to the non-normally distributed peak currents for the variant.

Here, we expressed either the wild-type α4 subunit or the variant α4(I114N) in HEK-293T cells in combination with β3 and δ subunits and compared their basal and GABA-activated function using whole-cell patch-clamp electrophysiology. The α4β3δ subunit combination was chosen as a background because it is the most abundant combination for α4-containing receptors (Choi et al., 2022; Sur et al., 1999). To assay for changes in GABA sensitivity, basal channel activity, and expression (estimated from peak current), cells were lifted into the flow of a multi-barrel perfusion tube and defined pulses of GABA and/or other ligands were delivered by translating the perfusion barrels using a piezoelectric actuator enabling sub-millisecond solution exchange. A microfluidic switch was used to swap solutions within one of the perfusion barrels to record a series of responses to a range of GABA concentrations for an individual cell (Fig. 1B). We observed similar GABA-sensitivity to that previously reported for wild-type α4β3δ receptors (Jiang et al., 2011; Patel et al., 2014). Peak current versus GABA concentration curves were fit with the Hill equation (Eq. 1). Although there is a slightly shallower curve in the α4(I114N)β3δ variant, the difference is small relative to the cell-to-cell variation (Fig. 1C). Thus, we conclude that the variant has similar GABA-sensitivity to wild-type.

Given the role of α4β3δ receptors in tonic extrasynaptic signaling (Nusser et al., 1998), we assessed unliganded opening in the absence of GABA by applying the pore blocker picrotoxin (PTX) which should block any basal conductance through spontaneously open channels (Feng & Forman, 2018; Wagner et al., 2005) (Fig. 1D). To compare current amplitudes across cells, we used the combination of saturating GABA and the allosteric activator propofol to estimate maximal activation of all channels (Fig. 1D). The much smaller current response to GABA as compared to the combination of GABA and propofol reflects the fact that GABA is a weak partial agonist at α4β3δ receptors as previously described (Feng & Forman, 2018; Meera et al., 2009). For each cell, we estimated the basal and GABA-elicited open probability by normalizing the current responses from zero in the presence of the pore blocker PTX to unity at the peak response to GABA and propofol combined (Fig. 1D). There were no observed differences in basal or GABA-elicited open probabilities between cells expressing α4β3δ or α4(I114N)β3δ receptors (Fig. 1E).

Across experiments, peak current responses to saturating GABA were overall reduced for the variant, with the median current across variant-expressing cells being half that of wild-type (Fig. 1F). We interpret this reduction in current amplitude as a reduction in the number of channels expressed in the plasma membrane. Although we cannot completely rule out the alternative possibility that the variant might reduce the single-channel conductance, the similar current responses and GABA-sensitivity to those of wild-type suggest that the variant has not grossly altered channel function. Despite an effect size indicating that variant-containing channels express only half as well as wild-type, the relatively large cell-to-cell variation in peak current suggests that this result should be taken as a cautious prediction. Further tests (preferably in neurons) are needed to better clarify changes to channel expression.

Although we did not observe any functional differences between α4β3δ and α4(I114N)β3δ receptors, we did not test for differential effects of other allosteric modulators such as benzodiazepines, Z-drugs, barbiturates, or neurosteroids. Another potential complication is that the stoichiometry of δ-containing receptors may be more varied than their γ-containing counterparts (Botzolakis et al., 2016; Kaur et al., 2009; Sente et al., 2022; Wongsamitkul et al., 2016), although our results are largely consistent with prior reports for α4β3δ receptors. In summary, we constrained our experiments to a basic functional characterization that does not find any obvious functional defects of the α4(I114N) variant, but suggests that the variant impairs channel expression which would cause a reduction in GABAergic tone that could contribute to the pathogenesis of epilepsy.

## Methods

### Plasmids

Human GABA_A_R subunit cDNAs for α4, α4(I114N), β3 and δ subunits were cloned into the pUNIV vector (Venkatachalan et al., 2007) and verified by sequencing of the entire subunit (Genscript). See Reagents for full sequences and Uniprot (UniProt Consortium, 2023) identifiers.

### Cell Culture and Transfection

HEK-293T cells (ATCC, CRL-3216) were cultured in MEM (Gibco, 11095-080) supplemented with 10% fetal bovine serum (Gibco, A5256701), 1% non-essential amino acids (Gibco, 11140-050), 1% sodium pyruvate (Gibco, 11360-070) and 1% penicillin/streptomycin (Gibco, 151140-122) at 37 °C and 5% CO_2_. For electrophysiological experiments, cells were plated on glass coverslips (Assistent, 92100100030) and transfected at ∼60–80% confluency using FuGENE 4k (Promega, E951A). Human GABA_A_R subunit cDNAs for α4, α4(I114N), β3 and δ subunits were co-transfected at a 1:1:1 ratio (α:β:δ) with a total of 1 μg of protein per 35 mm dish of cells. Recordings were performed 24-48 hours post-transfection.

### Electrophysiology

Lifted whole-cell recordings were obtained at room temperature using an Axopatch 200A amplifier (Axon Instruments), ITC-16 digitizer (HEKA), and WinWCP acquisition software (Strathclyde). Cells were voltage-clamped at -60 mV and recorded currents were lowpass filtered at 1 kHz. Patch pipettes (3-6 MΩ) were pulled (Sutter, P-97-4511) from borosilicate glass (Sutter, BF150-86-10) and filled with an internal solution containing (in mM): 10 CsCl, 10 HEPES, 10 EGTA, 2 MgATP, and 110 CsF (pH 7.3 with KOH, ∼300 mOsm adjusted with sucrose as needed). The external buffer solution contained (in mM): 130 NaCl, 4 KCl, 10 CaCl_2_, 1 MgCl_2_, 10 HEPES, and 5 glucose (pH 7.3 with NaOH, ∼300 mOsm adjusted with sucrose as needed). GABA, PTX, and propofol were dissolved in the external buffer solution and applied to cells via a multi-barrel perfusion system (Warner Instruments, SF-77B). Defined pulses of GABA or other ligands were applied using a computer controlled piezoelectric stepper to rapidly switch perfusion barrels between buffer and ligand. Solution flow was controlled with a microfluidic pump (Elveflow, OB1-MK4). To obtain concentration-response curves for individual cells, the concentration of GABA in one of the perfusion barrels was changed using a microfluidic switch (Elveflow, MUX-Flow). The time to fully replace the solution in the barrel was ∼30 sec as determined by the time course of the loss of current responses to periodic pulses in the ligand barrel after switching the ligand barrel solution from saturating GABA to buffer only. We routinely waited >1 min to ensure complete concentration exchange.

### Data Analysis

Current traces and concentration-response curves were analyzed using custom Python scripts and Prism 11 (GraphPad). GABA concentration-response curves were fit with the Hill equation:

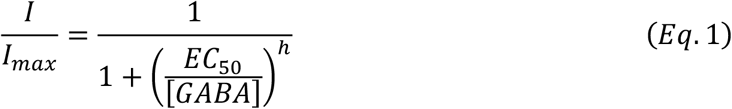

where *I* is the magnitude of the peak GABA-elicited current, *I*_*max*_ is the maximal peak response across GABA concentrations, [*GABA*] is the GABA concentration, *EC*_50_ is the concentration eliciting a half-maximal response, and *h* is the Hill slope.

Statistical comparisons were made using either a t-Test for normally distributed data or a non-parametric Kolmogorov-Smirnov t-Test for data distributions that deviate significantly from normal. A significance threshold of p < 0.05 was used for all analyses. Wild-type and variant experiments were run in parallel, and each dataset includes data from multiple experiment days except for the wild-type data in Fig. 1B-C which was collected in a single day. During the recording, data processing and analysis, group identity was not masked. No data were excluded from the reported datasets. Data are reported as mean ± SEM (symbol and error bars) or individual data and median ± interquartile intervals (symbols, line and box), as indicated. Box plots were generated in Python using the seaborn package.

## Reagents

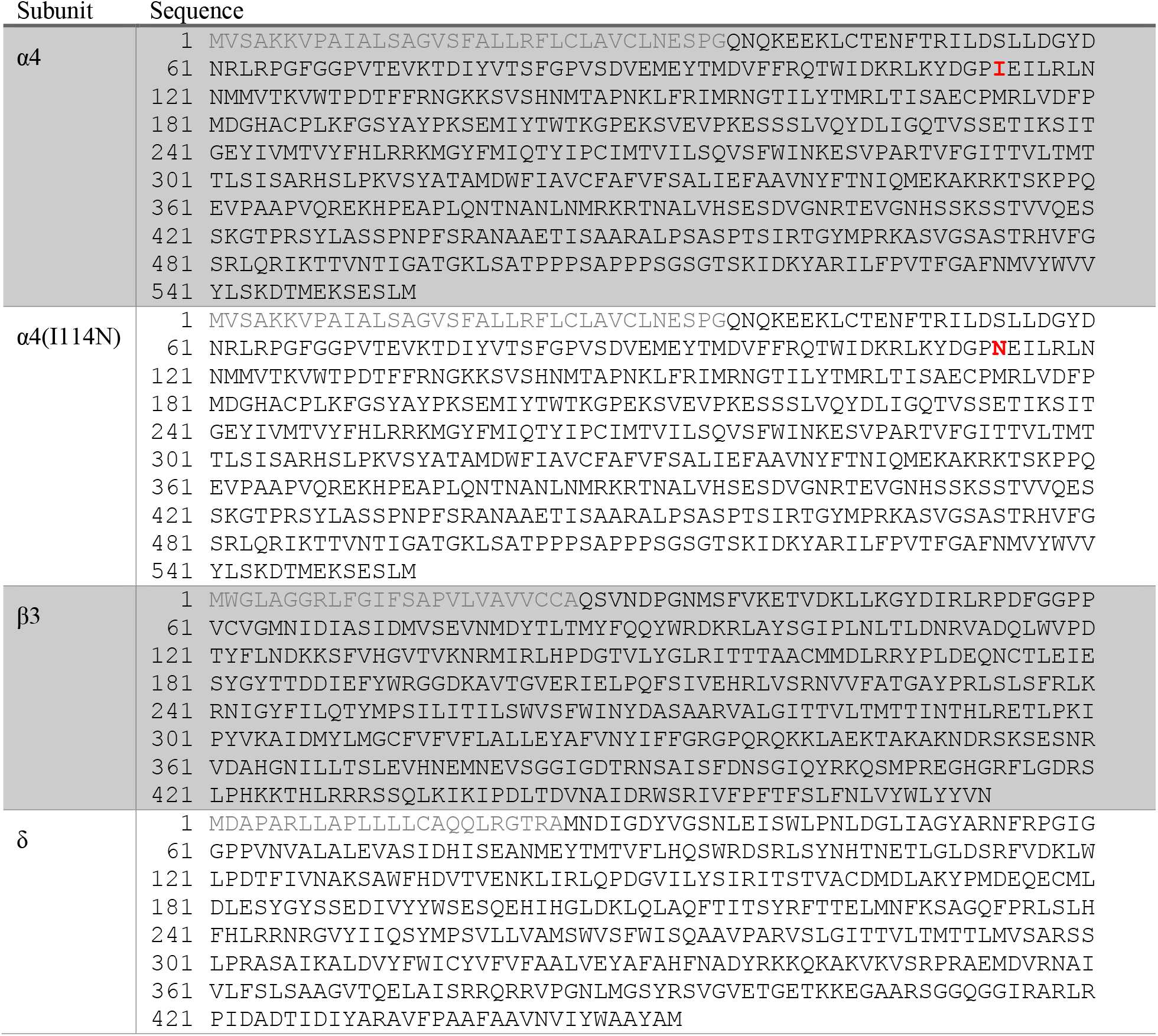

Uniprot sequence identifiers are α4: P48169, β3: P28472, and δ: O14764. Signal peptide shown in light gray. Position of α4(I114N) variant indicated in red/bold. Numbering starts from the first methionine in the signal peptide following HUGO nomenclature (Bruford et al., 2020). Using an alternative numbering convention often used in the scientific literature where numbering starts from the first residue in the mature protein following the signal peptide, the variant would be designated as α4(I79N).

## Acknowledgements

We sincerely thank the patient and his family for their participation in this study.

## Author Contributions

Conceptualization: M.P.G-O., A.C.B., and K.J. Data curation: D.W. and M.P.G-O. Formal analysis: D.W. and M.P.G-O. Funding acquisition: A.C.B. and M.P.G-O. Investigation: D.W., A.C.B., K.J., and C.M.B. Methodology (design of methodology): M.P.G-O. Project administration: M.P.G-O. Resources (Provision of study materials, reagents, materials, patients, laboratory samples, animals, instrumentation, computing resources, or other analysis tools.): A.C.B. and M.P.G-O. Supervision: M.P.G-O. Visualization: M.P.G-O., D.W. Writing — original draft: D.W., M.P.G-O., C.M.B., A.C.B., and K.J.

